# Transcriptional memories mediate the plasticity of sulfide stress responses to enable acclimation in *Urechis unicinctus*

**DOI:** 10.1101/2024.09.27.614670

**Authors:** Wenqing Zhang, Danwen Liu, Heran Yang, Tianya Yang, Zhifeng Zhang, Yubin Ma

## Abstract

To cope with environmental stresses, organisms often adopt a memory response upon primary stress exposure to facilitate a quicker and/or stronger reaction to recurring stresses. Somatic stress memory is essential in dealing with contemporary stress. The earliest sign of somatic stress memory is a change in gene transcription levels, which alters physiology and phenotype to better cope with stress. Sulfide is a common environmental pollutant; however, some organisms have successfully colonized sulfur-rich environments. Whether stress memory plays important role in sulfide stress adaptation remains unclear. In this study, to determine whether *Urechis unicinctus*, a sulfur-tolerant organism, retains the memory of previous sulfide stress, we simulated a repetitive sulfide stress/recovery system. The results showed that the tolerance of *U. unicinctus* to sulfide stress was significantly increased after priming with 50 µM sulfide. Further, transcriptional memory genes (TMGs) involved in regulating sulfide stress memory were identified, classified according to their expression patterns, and functionally analyzed. TMGs involved in sulfide metabolism, sugar metabolism, and protein homeostasis pathway showed an enhanced response, whereas those related to DNA repair pathway demonstrated a modified response pattern. Our study indicated that *U. unicinctus* retains memory of sulfide stress priming, which mediates plasticity to accelerate sulfide stress adaptation.

## 1. Introduction

During growth and development, organisms must cope with many biotic and abiotic repetitive stresses in nature, such as temperature, salinity, and pollutants, which require them to adapt to natural stresses (Oberkofler et al., 2021; Sharma et al., 2022; Tobler and Culumber, 2016). Stress memory is an important mechanism for stress adaptation (Zhang and Tian, 2022). Stress memory refers to the phenomenon in which organisms will have a higher tolerance when faced with same stress again after experiencing initial stress (Oberkofler et al., 2021). Stress memory can be used to cope with repetitive stress by modulating phenotypic changes in organisms. Corals are marine sessile organisms, many of which experience repeated marine heatwaves during growth, and stress primed corals exhibit a reduced incidence and/or severity of bleaching (Bay and Palumbi, 2015). This resilience is conferred by prior stress exposure and maintained through temporally varying stress events (Hackerott et al., 2021).

Antioxidant indicators reflect an organism’s ability to cope with environmental stresses, and many organisms typically possess a stress memory by regulating their antioxidant capacity when subjected to repeated and identical stresses after stress priming (Kambona et al., 2023). In response to repeated low-temperature stress, *Ciona robusta* relies on regulating the resilience of its antioxidant defence system (ADS) to maintain adaptive homeostasis in response to environmental challenges (Li et al., 2020). The mediterranean mussel (*Mytilus galloprovialis*) undergoes heat-hardening training, which results in the accumulation of amino acids associated with the regulation of osmosis and antioxidant. Heat stress alters the antioxidant metabolic profile of mussels, favoring increased heat tolerance in trained ones (Georgoulis et al., 2023). In addition, the threespine stickleback (*Gasterosteus aculeatus*) responds to repeated cold stress by increasing the expression of antioxidant genes (Kammer et al., 2011).

Most abiotic stresses are transient and repetitive, and contemporary organisms promote adaptation through somatic memory (Liu et al., 2022b). The most intuitive manifestation of somatic memory is changes in the transcriptional memory of genes, and these genes are called transcriptional memory genes (TMGs) (Avramova, 2015). TMGs can be classified into two type based on their mode of expression (Oberkofler et al., 2021). Type I TMGs exhibit sustained activation (or repression) in response to short-term stimuli; that is, genes that are persistently differentially expressed after the initial stress disappear. Genes that could be restored to non-stressed transcript levels after the disappearance of initial stress and were significantly differentially expressed (upregulation or downregulation) when experiencing the same stress again compared with initial stress were type II TMGs (Avramova, 2015; Oberkofler et al., 2021). TMGs are typically involved in the regulation of organismal homeostasis and play a cytoprotective role in the adaptive environmental memory of organisms (Ding et al., 2014; Friedrich et al., 2021; Galviz et al., 2020; Virlouvet et al., 2018). Type II TMGs are incredibly beneficial for sessile organisms that exposed to intricate environments, as they provide an efficient method for withstanding recurring environmental pressures (Hackerott et al., 2021). The enhanced and modified transcriptional responses exhibited by type II TMGs are equally important in biological responses to repeated stress (Gonzalez-Romero et al., 2017; Lim et al., 2021; Oberkofler et al., 2021). The expression levels of superoxide dismutase (*sod*) and peroxidase (*pox*) genes in wheat (*Triticum aestivum* L.) subjected to repeated high-temperature stress were significantly increased compared with the control plants without stress priming, suggesting that stress priming improves heat tolerance by enhancing the antioxidant capacity to mitigate the oxidative damage induced by high-temperature stress (Wang et al., 2014). In *Ciona robusta*, low-temperature stress priming rapidly adjusted the expression of genes in the oxidative stress response signaling pathway, and the magnitude of the response during repetitive stress was significantly weaker than that of the initial stress, suggesting that the organism remodeled the baseline of the antioxidant defense system after recovery from stress, resulting in a tolerant response (Li et al., 2020). Understanding stress priming induced transcriptional memory can provide a perspective for the generation of stress-tolerant organisms adapted to environmental change.

Among the many stressors, sulfide is one of the most special. Sulfide is the general name for H_2_S, HS^-^, and S^2-^, and a small amount of endogenous hydrogen sulfide is an important signal molecule in the organism, participating in a variety of physiological processes (Salehiyeh et al., 2024; Sun et al., 2022; Wu et al., 2019). However, excessive exogenous sulfide is a well-known environmental poison that must be removed through the sulfide metabolic pathway. At the micromolar level, it can cause metabolic damage and death (Evans, 1967; Nicholls, 1975). The mitochondria are the main sites of exogenous sulfide oxidation. Sulfide oxidation in mitochondria is mainly carried out by enzymatic reactions, which include four different but functionally related enzymes (sulfide: quinone oxidoreductase (SQR), persulfide dioxygenase (PDO), sulfite oxidase (SOX), and thiosulfate thiotransferase (TST)) that catalyse sulfide to thiosulfate and sulfate (Kabil and Banerjee, 2014; Paul et al., 2021). Sulfur-tolerant organisms show different physiological and biochemical response mechanisms to adapt to sulfide stress (Fujita et al., 2011; Greenway et al., 2020; Horsman et al., 2019; Kelley et al., 2016; Nobrega et al., 2024). The study of the molecular mechanisms of adaptation to sulfide is of great significance for explaining life phenomena in sulfide habitats (Greenway et al., 2020; Pfenninger et al., 2014; Rishi et al., 2024; Tobler et al., 2016; Wang et al., 2019).

Intertidal sediments are one of the main sites of sulfide enrichment, and sulfide concentrations in caves can reach 65 µM (Arp et al., 1992). *Urechis unicinctus* is a type of Echiura in the caves of coastal sediments that does not leave their caves for the majority of their entire life cycle and experiences repeated sulfide stress, which has a strong tolerance and detoxification ability to sulfide (Liu et al., 2015, 2022a; Ma et al., 2011, 2012; Zhang et al., 2013, 2021). Therefore, it can be considered as a model organism for studying sulfide metabolism and sulfide stress memory. Previous studies have shown a complete and efficient mechanism for sulfide oxidation (Li et al., 2018; Ma et al., 2011, 2012; Wang et al., 2023; Zhang et al., 2021). In *U. unicinctus*, the key enzyme of mitochondrial sulfide oxidation plays a significant role in sulfide stress adaptation, and it has an obvious stress response at the mRNA, protein, and enzyme activity levels, with the upregulated expression at the transcriptional level is the earliest and most obvious change (Liu et al., 2015; Ma et al., 2012). Furthermore, the involvement of DNA methylation in sulfide stress adaptation was identified by whole-genome bisulfite sequencing, RNA-seq and DNA methylase inhibitor validation in *U. unicinctus* (Zhang et al., 2024). Stress memory is an important mechanism for stress adaptation. However, it is not clear whether it plays an important role in sulfide stress adaptation. To determine the existence of sulfide stress memory and revealing its regulatory mechanisms that underlie sulfide stress memory are urgently required to enhance our understanding of sulfide stress adaptation.

In this study, we simulated a stress (S1)/recovery (R)/re-stress (S2) system by treating *U. unicinctus* with sulfide. First, by measuring survival and antioxidant indices in *U. unicinctus* after initial sulfide stress and repeated stress, we found repeated stress with 50 µM sulfide can trigger stress memory phenomenon. The expression patterns of TMGs were further characterized based on transcriptomic analysis during two rounds of sulfide stress. Combined with weighted gene co-expression network analysis (WGCNA) and kyoto encyclopedia of genes and genomes (KEGG) analyses, the key TMGs and pathways involved in sulfide stress memory were identified, providing new insights on adaptation to sulfide rich environments in organism.

## 2. Materials and methods

### 2.1 Animals and determination of survival

Adult *U. unicinctus*, measuring an average length of 13.5 ± 2.1 cm were collected from the coast of Yantai in Shandong Province, China. After acquisition, the worms were kept in aerated seawater (temperature: 20 °C, pH: 8.0, salinity: 30 PSU) for 3 days. Then, healthy worms were then randomly divided into groups of 12 worms each, and each group was kept in a sealed aquarium containing 30 L of seawater. Groups (each group with 3 biological replicates) were set up for the survival assay under 50 and 150 µM sulfide stress: the blank control group without exogenous sulfide stress, the control group not experiencing sulfide stress priming, and the stress priming group, in which the experimental individuals were stressed with 50/150 μM sulfide for 48 h, and then performed a 7 days recovery process. These groups were subjected to 50/150 µM sulfide stress treatment respectively, except for the blank control group. To ensure a realistic sulfide concentration, a sulfide stock solution comprising 10 mM Na_2_S and a pH level of 8.0 was added into sealed aquariums every 2 h, as measured using the methylene blue method (Cline and Richards, 1969; Arp et al., 1992). During the experimental period, dead individuals (worms that did not respond to mechanical stimulation) were removed, the number of dead in each group was recorded at 24 h intervals, and the survival rate was calculated.

### 2.2 Determination of antioxidant indices

The acquisition and culture of experimental animals and sulfide stress handling operations were partially the same as in Section 2.1. Then, 36 healthy worms were randomly assigned to 6 groups of 6 worms each, and each group was maintained in a sealed aquarium containing 30 L of seawater. The worms were initially exposed to 50 µM sulfide for 48 h (S1) and then transferred to natural seawater without exogenous sulfide for a 7 days recovery period, followed by another 48 h (S2) treatment with the same sulfide concentration. To control for potential confounding variables related to stocking density, 3 of the 6 aquariums were designated for sampling during the S1 period, whereas the remaining three aquariums were designated for sampling during the S2 period. 3 worms (one individual from each aquarium) were collected at 0, 6, 24, and 48 h during S1 and S2. A schematic of the treatment and sampling process is shown in Figure A.1. The collected hindgut tissues of worms were immediately immersed in liquid nitrogen to preserve their biological composition, and the frozen samples were stored at −80 °C. Antioxidant indices were determined for all samples, weighing 0.1 g of each sample (3 biological replicates), adding 9 times the volume of pre-cooled saline (0.9% NaCl), grinding it thoroughly in an ice bath, and centrifuging it for 10 min at 4 °C and 800 g. The supernatant was analyzed using CAT (A007), MDA (A003), T-AOC (A015), SOD (A001), and SOD (A015) test kits (Nanjing Jiancheng Bioengineering Institute, Nanjing, China) according to the manufacturer’s instructions. Data were analyzed using the PASW Statistics 18.0 (SPSS, USA) software package, and the significance of differences was tested by One-Way ANOVA, and two-by-two comparisons were made using Tukey’s method, with *P* <0.05 being considered a statistically significant difference.

### 2.3 RNA extraction, library construction and sequencing

First, RNA was extracted from the samples described in Section 2.2. Total RNAs from each stored tissue sample was extracted using TRIeasy™ LS Total RNA Extraction Reagent (Yeasen, Shanghai, China) following the manufacturer’s instructions. The yield, purity, and integrity of RNA samples were analyzed using a 2100 Bioanalyzer (Agilent Technologies, Santa Clara, CA, USA). A total of 24 sequencing libraries from 8 samples were generated using the Truseq^TM^ RNA sample prep kit (Illumina, USA). First, mRNA with poly (A) was purified from total RNA using oligo (dT) magnetic beads, fragmented, and reverse-transcribed into first-strand cDNA using random hexamer primers. Subsequently, second-strand cDNA was synthesized using RNase H and DNA polymerase I, end-repaired, and ligated with a sequencing adapter. The adaptor-ligated cDNA was PCR-amplified and sequenced on an Illumina NovaSeq 6000 platform.

### 2.4 Quality control and analysis of RNA-Seq

Sequencing data were analyzed for quality control using FastQC software, and reads were filtered to remove low-quality reads using Trimmomatic software. After obtaining clean reads, HISAT2 software (http://ccb.jhu.edu/software/hisat2/index.shtml) was used to compare the clean reads with the reference genomic data of *U. unicinctus*. Transcript assembly and quantification of each sample were completed using StringTie (http://ccb.jhu.edu/software/stringtie/) software. To determine the correlation between the sequenced samples, correlation analysis (pearson correlation analysis) and principal component analysis (PCA) were performed using normalized gene expression results. Differential expression analysis was performed on the normalized gene expression data using DESeq2, setting the criteria for differentially expressed genes (DEGs) as *P* <0.05 and |log_2_foldchange| >1 (Zhang et al., 2020). Co-expression networks were constructed using WGCNA for TMGs expression patterns. Genes that were not expressed in the tissues were removed prior to analysis. Soft thresholds were set based on a scale-free topology criterion. The correlation of each gene co-expression module with stress duration and number of stresses was assessed using the moduleTraitCor function to identify gene co-expression modules that were highly correlated with the traits. KEGG pathway enrichment analysis was performed on the set of DEGs using the clusterProfile software to identify the signaling pathways associated with the DEGs.

### 2.5 qRT-PCR analysis

The expression of genes in the 24 libraries was validated using quantitative real-time reverse transcription polymerase chain reaction (qRT-PCR). Total RNA isolation and qRT-PCR were performed as previously described (Chen et al., 2023). Three replicate analyses were conducted for each sample. The list of primers for the internal reference (*β-actin*) and other genes is shown in Table A.1. The cyclic threshold (Ct) was used for expression statistics, and the relative expression of target genes was expressed as 2^-ΔΔCt^. All data were expressed as mean ± standard error and analyzed statistically using SPSS software, with *P* <0.05 as a significant difference.

## 3. Result

### 3.1 Changes in survival and physiological indices of U. unicinctus after sulfide stress priming

To determine whether *U. unicinctus* was able to retain stress memory after sulfide stress priming, we established a stress-recovery-stress experimental system at two type of sulfide concentrations (50 μM and 150 μM). By comparing the survival rate of 50 µM (ecological concentrations) sulfide stress tolerance in the stress priming group and the control group, it was observed that individual deaths occurred in the control group on the 3rd day of stress, whereas in the stress priming group, individual deaths did not occur until the 6th day of stress. There were no individual deaths in the blank control group during the experiment period (Figure 1a). However, there was no significant difference in survival between the stress priming group and control group under the higher sulfide concentration (150 µM) stress (Figure 1b). We further determined the changes in antioxidant levels in the hindgut tissues under repeated sulfide stress with 50 µM sulfide. Malondialdehyde (MDA) content, total antioxidant capacity (T-AOC) activity, superoxide dismutase (SOD) activity, and catalase (CAT) activity can intuitively reflect the ability of organisms to cope with environmental stresses (Kambona et al., 2023; Li et al., 2020). We examined the content of these antioxidant indicators in samples from different treatment time points in the S1 and S2 periods. The results showed that the content of the peroxidation product MDA was reduced in the S2 period compared with the S1 period which showed a significant decrease at 6 h (Figure 1c). The T-AOC, SOD, and CAT activities were generally higher during the S2 period than during the S1 period (Figure 1d-f). In particular, T-AOC and CAT were significantly higher in samples treated for 6 h in S2 than in those treated for the same duration in S1 (Figure 1d, f). These data indicated that *U. unicinctus* subjected to stress priming with 50 µM sulfide could trigger stress memory when re-stressed with the same stress, as evidenced by an increase in tolerance time and a significant increase in antioxidant capacity.

**Figure 1.**
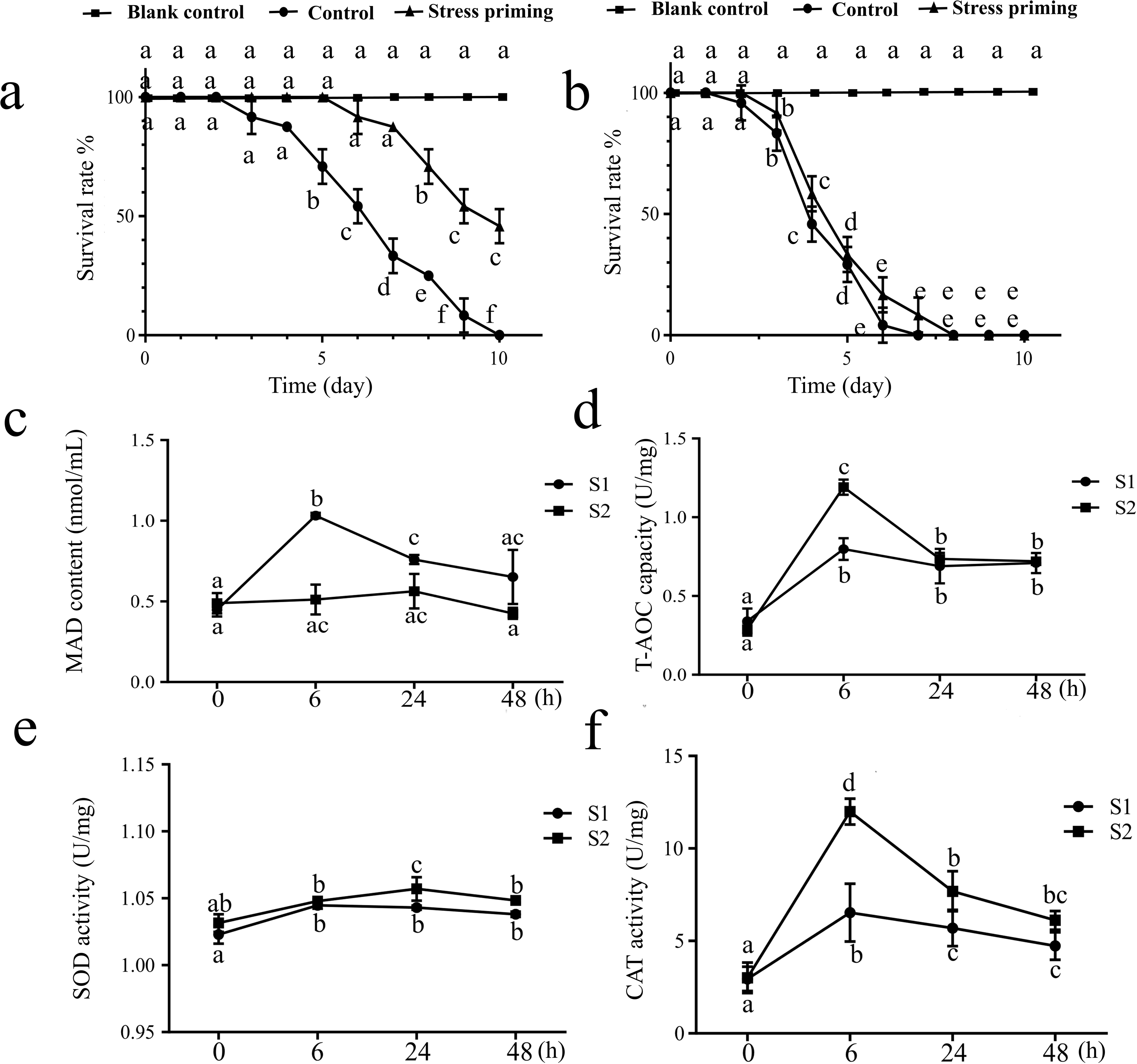
Survival of worms in different groups under 50 µM and 150 µM sulfide stress and determination of antioxidant indices in *U. unicinctus* under first stress (S1) and second stress (S2) with 50 µM sulfide. (a) Survival statistics of the stress priming group experiencing 50 µM sulfide stress priming, the control group not experiencing 50 µM sulfide stress priming, and the blank control group without sulfide stress, during the 50 µM sulfide stress process. (b) Survival statistics of the stress priming group experiencing 150 µM sulfide stress priming, the control group not experiencing 150 µM sulfide stress priming, and the blank control group without sulfide stress, during the 150 µM sulfide stress process. (c) Determination of malondialdehyde (MDA) content during S1 and S2. (d) Determination of total antioxidant capacity (T-AOC) activity during S1 and S2. (e) Determination of superoxide dismutase (SOD) activity during S1 and S2. (f) Determination of catalase (CAT) activity during S1 and S2. Different letters indicate significant differences between the data (*P* <0.05).

### 3.2 Analysis of transcriptome data and identification of TMGs under two rounds of sulfide stress

To characterize the TMGs involved in sulfide stress memory, we further analyzed the transcripts of the two rounds of sulfide stress. A total of 24 samples (3 biological replicates per sample) were sequenced at 4 time points (0 h, 6 h, 24 h, 48 h) in S1 and S2, respectively. One billion forty-three million raw reads and one billion sixteen million clean reads were generated, with a total sequencing volume of 162.4 G. Q20 ≥97.57% and Q30 ≥93.83%, the library quality met the requirements for subsequent analysis (Table A.2). The obtained clean reads were alignment with the *U. unicinctus* genome, and total mapped rate (%) was 83.78%, and the unique mapped rate (%) was 74.90% (Table A.3). The pearson correlation coefficient, which was calculated based on the expression value of each library, revealed a strong correlation between sample replicates (Figure A.2a). PCA of the samples further confirmed the consistency between replicates and the gradual differentiation of samples at the same time points in S1 and S2 as the duration of stress increased (Figure A.2b).

We counted the DEGs (|log_2_FoldChange| >1, *P* <0.05) between S1_6, S1_24, S1_48 and the control group (S1_0) samples in S1, respectively, and the same analysis was performed in S2. The analysis showed that *U. unicinctus* produced a large number of DEGs during both S1 and S2. The numbers of DEGs produced during S1 being 2004 (S1_6 vs S1_0), 3185 (S1_24 vs S1_0) and 4392 (S1_48 vs S1_0) (Figure 2a). The numbers of DEGs produced during S2 were 2032 (S2_6 vs S2_0), 2496 (S2_24 vs S2_0) and 7230 (S2_48 vs S2_0) (Figure 2b). The number of DEGs showed a time-dependent increase during sulfide stress, and the number of DEGs was significantly higher in S2 than in S1 (Figure 2a-b). We further counted the DEGs between the samples at the same time points of S1 and S2, and the number of DEGs was 2189 (S2_0 vs S1_0), 1937 (S2_6 vs S1_6), and 2273 (S2_24 vs S1_24) 4648 (S2_48 vs S1_48) (Figure 2c). After filtering out the duplicated differential genes, a total of 7804 DEGs were obtained (Figure 2d). According to the definition of memory genes (Avramova, 2015), the above 7804 genes can be regarded as sulfide stress TMGs. To further validate the credibility of the transcriptome data, we selected among these TMGs the key enzymes *sqr* and alternative oxidase (*aox*) involved in sulfide metabolism, as well as the antioxidant-related enzymes *sod* and catalase (*cat*), which were genetically quantified using qRT-PCR in sequenced samples. The results showed that the patterns of these genes were consistent with the trends in gene expression changes in RNA-Seq (Figure A.3). The above genes were significantly upregulated at each time point of sulfide stress in S2 compared with S1, and the faster and stronger response contributed to accelerate sulfide metabolism and removed reactive oxygen components, thus improving the sulfide stress tolerance ability of *U. unicinctus*.

**Figure 2.**
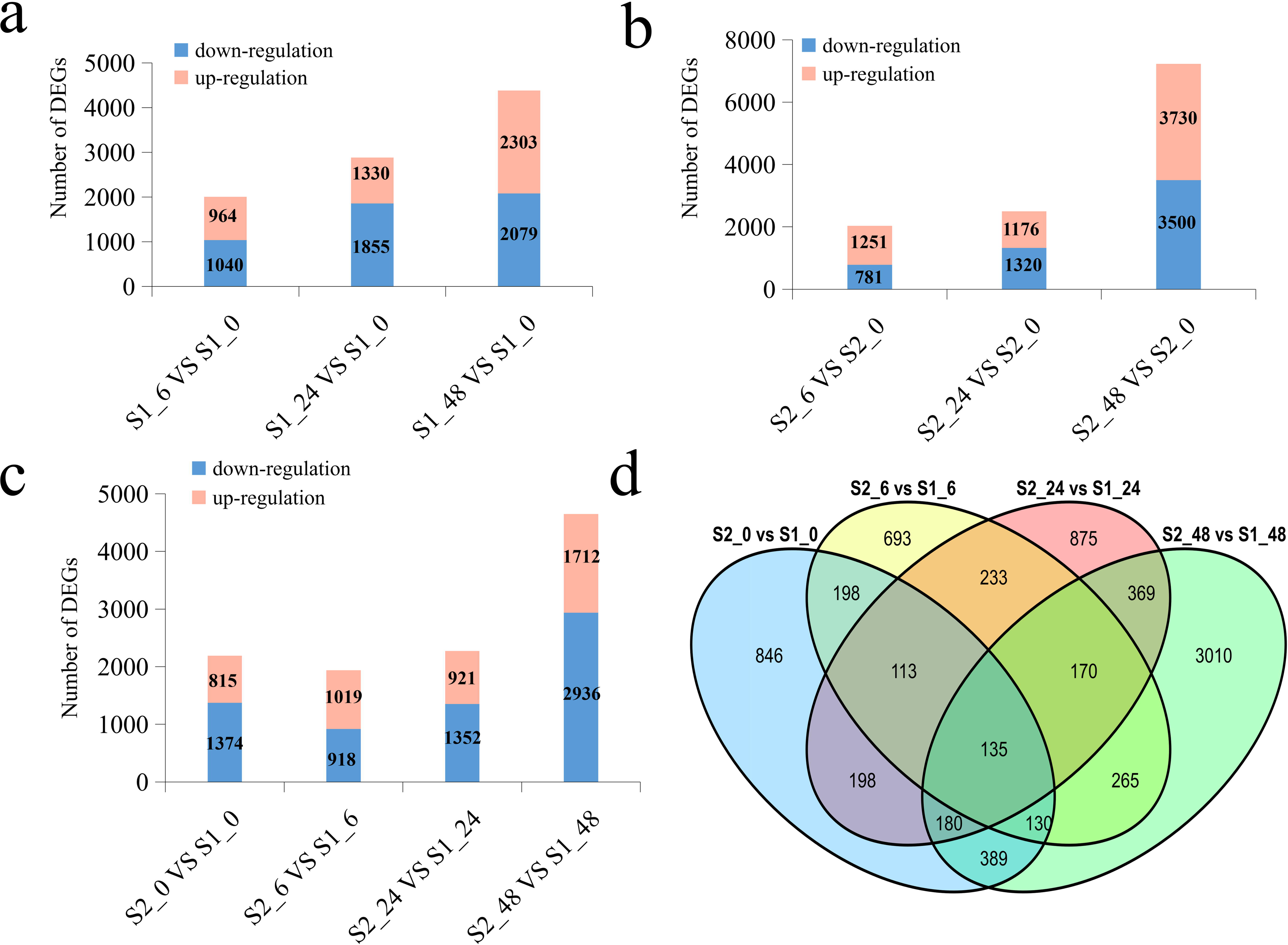
Statistical analysis of DEGs of *U. unicinctus* after repeated sulfide stress. (a) Statistical analysis of DEGs in S1. (b) Statistical analysis of DEGs in S2. (b) DEGs between the same time points in S1 and S2. (d) Venn diagram of DEGs in S1 and S2.

### 3.3 Expression pattern analysis of TMGs

Modular analysis of the transcriptomic data association by WGCNA was used to identify the expression pattern of TMGs. First, to improve the accuracy of co-expression network construction, low-quality TMGs (FPKM <1, coefficient of variation <0.5) were filtered out, and 5572 TMGs obtained from the screening were used for analysis. Then, we performed WGCNA analysis of the above TMGs. The 5572 TMGs were divided into 14 gene co-expression modules, with the highest number of TMGs (2320) in module 8; the lowest number (52) in module 9. The module 14 which represents a collection of genes that could not be assigned to any of the modules and was not analyzed subsequently (Figure 3ab).

**Figure 3.**
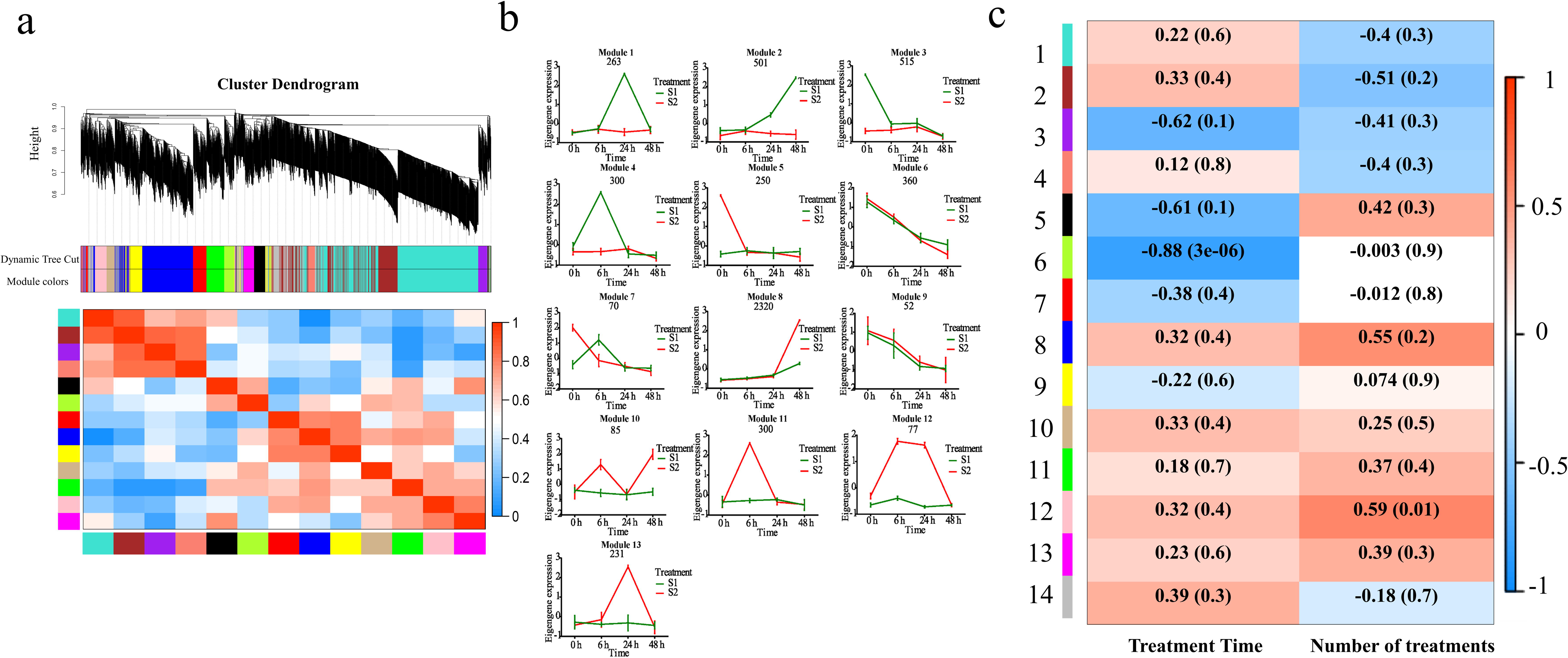
WGCNA analysis of TMGs, expression patterns of characteristic genes in different modules and heat map of correlation analysis between gene co-expression modules and traits. (a) WGCNA analysis of TMGs. (b) Expression patterns of characteristic genes in 13 modules. (c) Heat map of correlation analysis between gene co-expression modules and traits during repeated sulfide stress process. Each row corresponds to a gene co-expression module (different modules specified by WGCNA are numbered 1-14), and each column represents a trait, displaying the correlation between the modules and sulfide stress time and the number of sulfide stress treatments; the numerical values in each grid represent the correlation coefficient between the module and the trait, as well as the corresponding *P*-value (in parentheses); the red and blue squares represent positive and negative correlations between traits and modules, respectively.

By analyzing the gene expression profiles of the genes in the 13 modules in S1 and S2, we found that the TMGs in modules 3, 5, and 7 belonged to type I TMGs (significantly differentially expressed between S2_0 and S1_0). The TMGs in modules 1, 2, 4, 8, 10, 11, 12, and 13 belonged to type II TMGs (significantly differentially expressed between S2_n and S1_n; n = 6, 24, or 48), in which the genes in modules 8, 10, 11, 12, and 13 had an enhanced magnitude of response in S2 compared with S1 and were defined as enhanced response TMGs modules. The genes in modules 1, 2, and 4, which had an attenuated magnitude of response in S2 compared with S1, were defined as modified response TMGs modules (Ding et al., 2014). Overall, Type I TMGs totaled 835. There was a total of 4077 type II TMGs, of which 3013 belonged to the enhanced response and 1064 to the modified response (Figure 3c).

### 3.4 Identification of key pathways and TMGs involved in sulfide stress memory

Furthermore, we analyzed the correlation of the above gene modules with sulfide stress time (0, 6, 24, and 48) and the number of sulfide stress treatments (1 and 2). We found that modules 2, 8, and 12 had the highest correlation with the number of sulfide stress treatments, with correlation coefficients >0.5, and the genes in these modules were highly correlated with the sulfide stress memory phenomenon (Figure 3c). The expression of TMGs in module 2 was consistently upregulated during the S1 period; however, the expression of the above genes was downregulated at the 24 h and 48 h of S2 compared with the S1 period (Figure 3b; Table A.4). Such modified response TMGs are usually involved in the regulation of the homeostatic restoration of organisms in response to repeated stress. KEGG pathway enrichment analysis showed that the genes in this module were mainly enriched in immune stress signaling pathways such as amino acid synthesis and metabolism, TNF signaling pathway, and toll-like receptor signaling pathway (Figure 4a). We further analyzed the functions of the top10 core genes in this module based on the eigengene connectivity (kME) value, which mainly includes ATP-dependent DNA helicase Q1 (*RecQ1*), replication protein A (*RPA*), histone N-methyltransferase SUV420H1 (*SUV420H1*), and other genes involved in DNA repair reaction-related genes (Table A.5).

**Figure 4.**
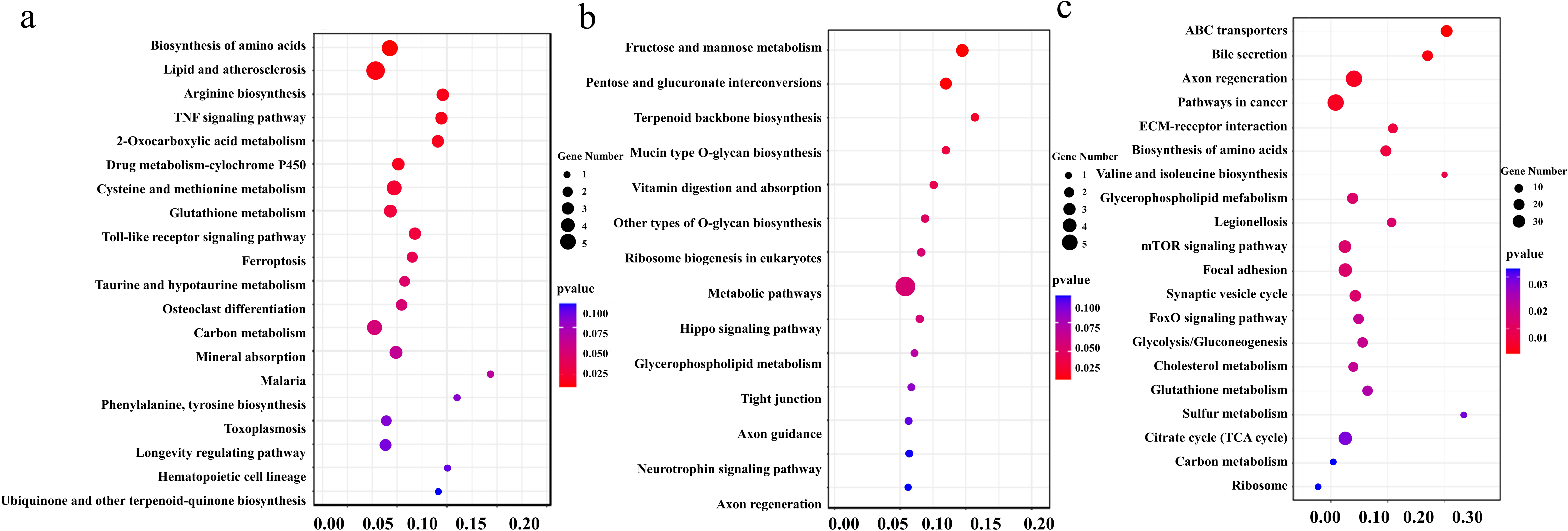
KEGG enrichment analysis of TMGs in representative modules. (a) KEGG enrichment analysis of TMGs in module 2. (b) KEGG enrichment analysis of TMGs in module 12. (c) KEGG enrichment analysis of TMGs in module 8.

Compared with S1 period, TMGs were significantly upregulated in module 8 and module 12, during S2 period (Figure 3b). Genes with this type of expression pattern are enhanced response TMGs, which typically play a cytoprotective role in organisms. The genes in module 12 showed the above expression patterns at 6 and 24 h (Figure 3b; Table A.4). KEGG enrichment analysis showed that the genes in this module were mainly concentrated in sugar metabolism pathways such as fructose and mannose metabolism (Figure 4b), and the top10 core genes in this module were c-type lectin, ras-related protein Rab-21, carbohydrate sulfotransferase 1, and other related genes involved in substance transportation, detoxification, and immunity (Table A.6).

Module 8 is the module with the highest number of TMGs, totaling 2320. Compared with module 12, it showed an enhanced response later which showed an enhanced response at 48 h (Figure 3b; Table A.4). KEGG enrichment analysis showed that the genes in this module were mainly enriched in sulfide metabolism, transporter protein-mediated substance transport processes, mTOR and FoxO signaling pathways, and glycolysis/gluconeogenesis (Figure 4c). Notably, the mitochondrial oxidative pathway, which plays an important role in sulfide detoxification, contains four key enzymes, and three of the key enzyme genes (*sqr*, *tst*, and *pdo*) were significantly enriched in the sulfide metabolic pathway. The transcript levels of these genes showed an enhanced response during the second sulfide stress (Figure A.4). The significantly enriched transporter proteins were mainly from two protein families, ATP-binding cassette transporter proteins (ABC) and solute carrier proteins (SLC) in the bile secretion pathway. In addition, the heat shock protein 70 (HSP 70) family, which is involved in the intracellular oxidative stress pathway, as well as *FoxO*, a core transcription factor in the FoxO signaling pathway, is also among the top 10 core genes in this module (Table A.7). Taken together, it is possible that this module is involved in the late response to repeated sulfide stress.

## 4. Discussion

### 4.1 Stress memory occurs during repetitive sulfide stress in U. unicinctus

Fluctuating environmental changes place enormous stress on organisms in marine ecosystems (Lim et al., 2021). Organisms previously exposed to stress may alter their responses to subsequent stresses, thereby remodeling the organism’s physiological responses and phenotypes, and triggering stress memories to cope with the complex external environment (Ding et al., 2014). Annual changes in ocean heat waves (1982-2017) and coral bleaching rates in 14 regions of the Great Barrier Reef, Australia, showed that repeated ocean heat waves have resulted in significant reductions in bleaching rates of *Porites*, suggesting triggered stress memory (DeCarlo et al., 2019; Hackerott et al., 2021). In *Daphnia magna*, researchers have found that parental (F0) experiencing salt stress can lead to increased fertility in its first generation (F1), which can be sustained to the third generation (F3). High salt stress can induce stress memory in *Daphnia magna*, which can be inherited across generations (Jeremias et al., 2018).

Exogenous sulfide is a common pollutant in nature and can affect the normal development and growth of organisms at micromolar levels (Chou et al., 2023). Examples of gradual biological adaptations to sulfide exposure have been reported. Fish of the same lineage (*Poecilia mexicana*) show different phenotypes and gene expression patterns depending on the sulfide concentration in their habitat (Kelley et al., 2016). Further studies have shown that DNA methylation sites specific to the fish adapted to survive in sulfide can be inherited across generations (Kelley et al., 2021). However, whether repeated sulfide stress triggers the stress memory mechanism in organisms, and the key pathways and genes involved in sulfide stress memory remain unclear.

To determine whether *U. unicinctus*, which recognized as sulfur-tolerant organism, retained memory of priming sulfide stress, we designed a repetitive stress/recovery system for its ecological and higher concentrations. We found that in *U. unicinctus*, under near-ecological concentrations (50 µM) of sulfide stress, the semi-lethal time was significantly longer in the stress priming group. However, after experiencing stress priming with 150 µM sulfide, which far exceeded the ecological concentration, *U. unicinctus* did not show improved tolerance to secondary stress. This is consistent with previous findings that stress responses within the acceptable range of organisms are capable of inducing the phenomenon of stress memory (Ding et al., 2012; Song et al., 2024). The response of organisms to environmental stress depends on the intensity and duration of the stress (Hackerott et al., 2021). Thus, the stress priming dose mediates the intensity of the response to subsequent stresses. If the duration or intensity of the initial initiating exposure is insufficient, or both, a detectable memory response may not be triggered, whereas exposures exceeding tolerable thresholds can weaken the organism to the point where any potential benefit is lost (Hackerott et al., 2021). Our study showed that *U. unicinctus* retains memory of 50 µM sulfide stress priming, which in turn mediates plasticity to accelerate adaptation to repeated sulfide stress. The sulfide stress memory system of *U. unicinctus* developed in this study is an important guide for organisms living in the same environment to cope with sulfide stress better.

The increase in stress tolerance may be closely related to changes in antioxidant levels (Kambona et al., 2023). Dynamic regulation of antioxidant indices is a marker of changes in biological stress capacity, and organisms typically possess stress memory by remodeling their antioxidant baselines (Li et al., 2020). To determine whether sulfide stress priming reshaped the antioxidant baseline, we further measured the antioxidant indices in response to 50 µM sulfide stress after stress priming in *U. unicinctus*. Among them, the accumulation of MDA may cause some damage to membranes and cells, so its content can reflect the degree of adverse damage suffered by organisms (Del Piano et al., 2024). The biological function of CAT and SOD are to promote the decomposition of H_2_O_2_ in cells so that it does not produce further hydroxyl radicals with high toxicity, and thus protects the functional role of the antioxidant system (Gautam et al., 2006). T-AOC can reflect the total antioxidant level of the endosomes of organisms (Jia et al., 2012). In our study, MDA content in *U. unicinctus* was significantly reduced in response to repeated sulfide stress, indicating that stress priming could reduce the damage during repeated stress, whereas CAT, SOD and T-AOC activities were significantly elevated, suggesting that *U. unicinctus* improved their ability to cope with stress after priming. These results showed that sulfide stress priming significantly increased the sulfide tolerance by remodeling the antioxidant baseline, thus inducing stress memory. These findings have important implications for organisms experiencing sulfide stress in the environment.

### 4.2 Identification of key TMGs and pathways involved in sulfide stress memory

Stress memory mechanism is an important means by which organisms adapt to complex environments (Charng et al., 2023; Demongeot et al., 2019). Somatic stress memory plays an important role in coping with contemporary repetitive stress (Liu et al., 2022b). The most intuitive and preliminary manifestation of somatic stress memory is changes in the level of gene transcription in response to repetitive stress, and these TMGs can reshape the physiology and phenotype of the organism to better cope with repetitive stress (Ding et al., 2012). Our analysis of transcriptome data from two rounds of sulfide stress revealed that *U. unicinctus* have the same two types of TMGs that have been demonstrated in other organisms (Ding et al., 2014). TMGs with the same expression trend were grouped together, and modules closely related to sulfide stress treatment were obtained by WGCNA analysis. Further combined with KEGG enrichment analysis, the key TMGs and pathways involved in sulfide stress memory were identified. Finally, we obtained a map of the regulatory network of genes involved in sulfide stress memory in *U. unicinctus* (Figure 5). In response to repeated sulfide stress, *U. unicinctus* enhanced cellular protection by enabling enhanced responses of TMGs in pathways, such as sulfide metabolism, antioxidant enzyme system, energy metabolism, and maintenance of protein homeostasis. Simultaneously, the modified response of TMGs involved in DNA repair may lead to a decrease in sensitivity and an increase in tolerance to repeated sulfide stress.

**Figure 5.**
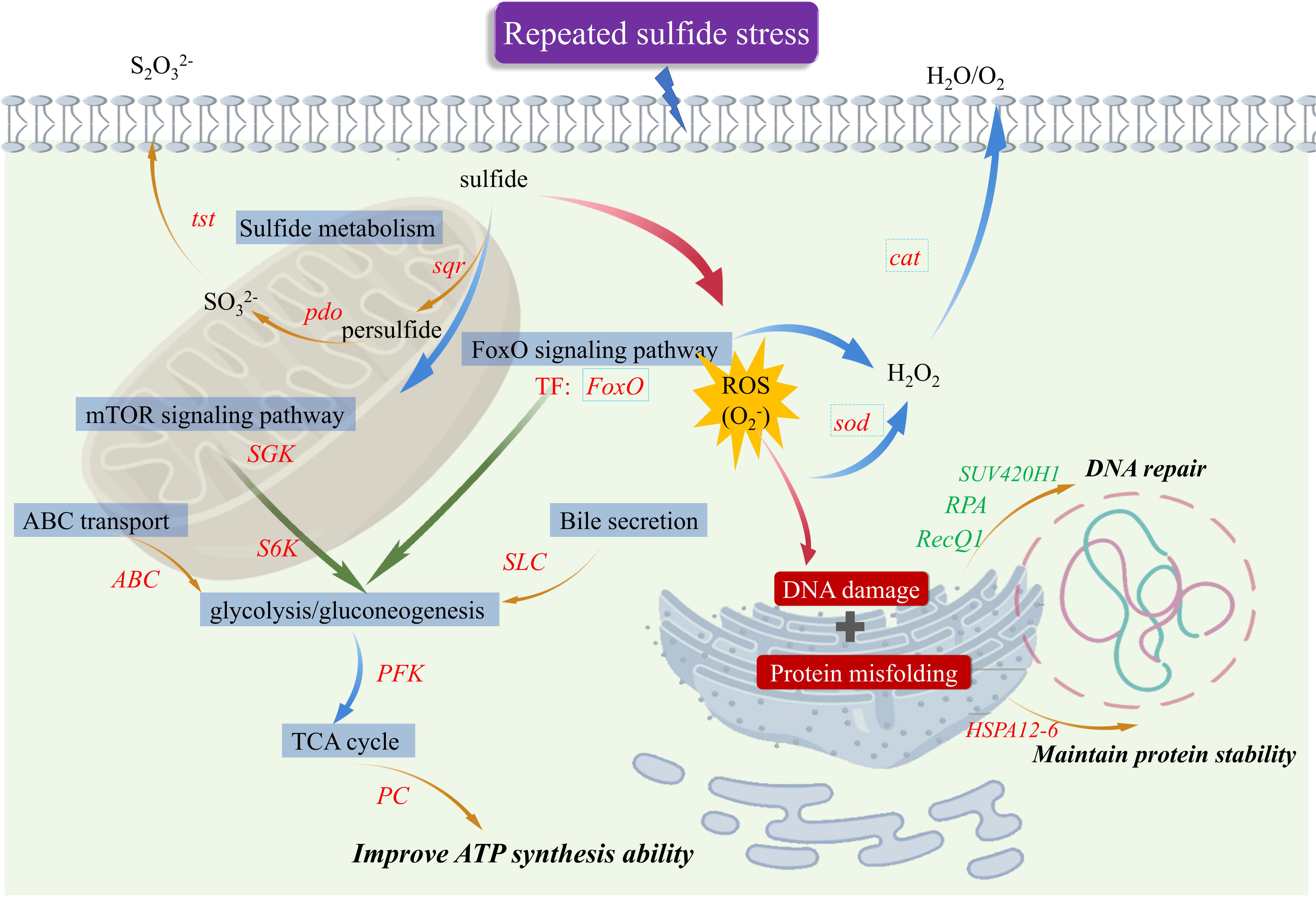
The regulatory network diagram of TMGs under repeated sulfide stress in *U. unicinctus.* Red font represents enhanced response TMGs, green font represents modified response TMGs. *tst*, thiosulfate thiotransferase; *pdo*, persulfide dioxygenase; *sqr*, sulfide: quinone oxidoreductase; *SGK*, glucocorticoid-regulated kinase; *S6K*, ribosomal protein S6 kinase; *ABC*, ATP-binding cassette; *SLC*, solute carrier; *PFK*, phosphofructokinase; *PC*, pyruvate carboxylase; *sod*, superoxide dismutase; *cat*, catalase; *SUV420H1*, histone N-methyltransferase SUV420H1; *RPA*, replication protein A; *RecQ1*, ATP-dependent DNA helicase Q1; *HSPA12-6*, Heat Shock Protein A12-6; SO_3_ ^2-^, sulfite; S_2_O_3_ ^2-^, thiosulfate; ROS, reactive oxygen species; TF, transcription factor. The genes within the solid box are transcription factors, whereas those within the dashed box represent their downstream target genes.

#### 4.2.1 Sulfide metabolism-associated TMGs respond to repeated sulfide stress by enhanced response patterns

The invasion of exogenous sulfide into an organism results in the impairment of a wide range of life activities (Wei et al., 2021; Zhao et al., 2023). However, some sulfur-tolerant organisms can reduce or even relieve the toxicity of sulfide invading the organism and its damage through oxidative sulfide metabolic pathways (Ma et al., 2012; Tiranti and Zeviani, 2013; Zhang et al., 2024). The mitochondria are the main sites for sulfide oxidation (Chen et al., 2021). Sulfide oxidation in the mitochondria is mainly carried out by enzymatic reactions, which include four different but functionally related enzymes that work together to catalyse sulfide to thiosulfate and sulfate (Ma et al., 2011; Thakur and Anand, 2021). Sulfide is oxidised in the mitochondria by SQR to generate persulfide, which is the first step in catalysis of mitochondrial oxidation. It is then further oxidised to sulfite (SO_3_^2-^) by PDO. SO_3_ ^2-^ can be oxidised to sulfite (SO ^2-^) by SOX and reduced to thiosulfate (S O ^2-^) by TST (Kabil and Banerjee, 2014; Paul et al., 2021). Some sulfur-tolerant organisms have been found to respond to sulfide stress by increasing the expression levels of key genes in the mitochondrial oxidation pathway. For example, the highly sulfur-tolerant razor clam (*Sinonovacula constricta*) relies on the mitochondrial oxidative pathway to detoxify sulfide, with the *sqr* gene being the most responsive at the transcriptional level (Chen et al., 2021). The *Gigantidas platifrons*, known as deep-sea mussel, can survive in H_2_S-rich environments such as hydrothermal vents. It has been demonstrated that deep-sea mussels adapt to H_2_S-rich environments mainly by upregulating the transcript levels of the genes related to oxidative phosphorylation and mitochondrial sulfide oxidation pathway (Sun et al., 2022).

However, excess exogenous sulfide activates oxidative stress pathways, which can cause cellular damage (Wang et al., 2012; Zhao et al., 2023). The FoxO signaling pathway is involved in the regulation of many cellular physiological events, including anti-oxidative stress, glucose metabolism, apoptosis, and lifespan extension (Guo et al., 2024; Link, 2019; Putker et al., 2013; Song et al., 2023). The antioxidant enzyme genes *sod* and *cat* have been shown to be downstream targets of the transcription factor *FoxO*, and upregulation of their expression can alleviate cellular damage caused by oxidative stress (Donate-Correa et al., 2023; Zečić and Braeckman et al., 2020). In this study, three key genes of the mitochondrial oxidative pathways (*sqr*, *tst*, and *pdo*) were classified into module 8, showing an enhanced response pattern of transcriptional memory. Meanwhile, the FoxO signaling pathway, which is involved in anti-oxidative stress, was significantly enriched in Module 8. Among them, *FoxO* was the top10 core gene of this module, indicating *FoxO* may be the key molecule involved in the memory of sulfide stress in *U. unicinctus*. Thus, the initial stress triggered a stress memory response in the expression of key enzyme genes for sulfide metabolism and anti-oxidative stress, and they improved the efficiency of oxidative detoxification of sulfide through a stronger upregulation of gene expression response.

#### 4.2.2 DNA repair-associated TMGs respond to repeated sulfide stress by modified response patterns

DNA is the carrier of genetic information, and the maintenance of normal transcription of genes is a prerequisite for the implementation of various physiological activities in living organisms and is essential for normal life processes and species continuation (Abdelazeem et al., 2021; Shackelford et al., 2021). Organisms may accumulate reactive oxygen species (ROS) due to biotic or abiotic stress, and when they accumulate excessively, they cause DNA damage, which is mainly manifested by double-stranded or single-stranded DNA breaks, leading to the normal life activities of the organisms being affected or even death (Abdolsamadi et al., 2020; Abdulazeez et al., 2022). Previous studies have reported that sulfide induces the formation of ROS in organisms (Shackelford et al., 2021). This is one of the reasons why many organisms can not survive in sulfur-rich environments (Fu et al., 2018; Shackelford et al., 2021). Therefore, increased tolerance to DNA damage is an important mechanism to maintain the integrity of the cellular genome and the normal survival of the organism in case of excessive accumulation of ROS (Fu et al., 2018). During DNA repair, the *SUV420H1* gene, which is associated with chromatin modification, recruits repair factors and participates in the DNA damage repair process (Bröhm et al., 2019). RPA is a DNA repair-associated protein that accomplishes the detection and repair at DNA damage sites together with a number of proteins responsible for chromosome structure maintenance, protection, and repair functions (Oakley and Patrick, 2010). RecQ1, on the other hand, is a multifunctional DNA-deconjugating enzyme that interacts with DNA repair proteins, such as RPA, to repair DNA damage (Debnath and Sharma, 2020). During repetitive sulfide stress in *U. unicinctus*, DNA repair-related TMGs (*RecQ1*, *RPA*, *SUV420H1*) presenting a modified response pattern were classified into module 2 and were the top10 core genes. These results indicate that genes in the DNA repair pathway that respond to decreased sensitivity to repeated sulfide stress may contribute to the remodeling of the defense baseline, causing increased sulfide tolerance in *U. unicinctus*.

#### 4.2.3 Maintenance of protein homeostasis-associated TMGs respond to repeated sulfide stress by enhanced response patterns

HSP70 performs the most basic physiological functions in cells, such as protein folding, stretching, transport, oligomer formation, and depolymerization, to maintain cell survival and function, and overexpression of HSP70 plays a stress-protective role by maintaining protein homeostasis under unfavorable conditions of stress (Chan and Groisman, 2024; Sojka et al., 2024). It has been found that *HSP70* plays an active role in the stress response to a variety of biotic and abiotic factors (Hu et al., 2019; Li et al., 2024; Wang et al., 2020). Expression of HSP70 family members was significantly upregulated in *Ruditapes decussatus* when subjected to repeated infestation with *Micrococcus luteus* (El-Wazzan et al., 2019). Transcript levels of HSP70 family members were significantly upregulated in the progeny of *Artemia franciscana* infected by *Vibrio campbellii* when their reproducing offspring were re-infected compared with the offspring produced by the uninfected parent (Norouzitallab et al., 2016). HSPA12-6 is a member of the HSP70 family, which plays an important role in the maintenance of protein homeostasis and regulation of biological processes in cells (Linder and Pogge von Strandmann, 2021; Liu et al., 2022a). In this study, multiple HSP70 family genes (*HSPA12-6, HSP70b2-1*, and *HSP70b2-2*) were categorized in module 8. Among them, *HSPA12-6* was the top10 core genes of this module. *U. unicinctus* experiencing repetitive sulfide stress might enhanced transcriptional memory of the above TMGs related to maintaining protein homeostasis, a means to better cope with repetitive sulfide stress.

#### 4.2.4 Sugar metabolism-associated TMGs respond to repeated sulfide stress by enhanced response patterns

Sugar metabolism is an important energy-supplying pathway in the biological response to environmental stress (Li et al., 2017; Mizock, 1995; Oláh et al., 2008). Under aerobic conditions, pyruvate, an important intermediate product of sugar metabolism, enters the mitochondria to produce acetyl coenzyme A, which is further oxidized via the tricarboxylic acid (TCA) cycle to produce ATP, water, and carbon dioxide (Somsen et al., 2000; Wang et al., 2024). However, under hypoxic conditions, pyruvate is converted to lactate and produces ATP via the glycolysis pathway, in which phosphofructokinase (PFK) and pyruvate carboxylase (PC) are the key enzymes in the glycolytic process (Kresge et al., 2005; Speers-Roesch et al., 2013; Yin et al., 2009). In addition, several signaling pathways are involved in regulating glucose metabolism. The mTOR and FoxO signaling pathways sense changes in various intracellular signals, including growth factors, hormones, nutrient factors (glucose and amino acids), and stress conditions (Cheng et al., 2014; González et al., 2023). When activated, the mTOR signaling pathway can be used to mediate the glycolytic pathway and the TCA cycle by glucocorticoid-regulated kinase (SGK) and ribosomal protein S6 kinase (S6K) (Cheng et al., 2014). ABC in the ABC transporter pathway and SLC in the bile secretion pathway, on the other hand, are mainly involved in sugar metabolism through the glucose transporter pathway (Choudhuri and Klaassen, 2006; Morioka et al., 2018). Excess sulfide blocks electron transfer in the respiratory chain, inhibiting aerobic respiration, thus, affecting the efficiency of energy production (Shen et al., 2020). It was found that the expression levels of genes involved in the glycolytic pathway were significantly higher in the fish (*Poecilia mexicana*) adapted to survive in sulfide-rich environments than in their sulfide-free counterparts, and that the sulfide-tolerant fish compensated for the lack of energy produced by aerobic respiration owing to excess sulfide by accelerating the glycolytic pathway in their organisms (Kelley et al., 2016). In module 8, TMGs were significantly enriched in glycolysis/gluconeogenesis and TCA cycle metabolic pathways. Meanwhile, the mTOR signaling pathway, ABC transport and bile secretion pathway, which are involved in the regulation of glucose metabolic pathways, were also significantly enriched. The significant enrichment of TMGs in sugar-based metabolic pathways is believed to be involved in the regulation of energy metabolism to alleviate the adverse effects of sulfide stress on normal physiological activities, which is of positive significance for effective defense against repeated sulfide stress. This suggests that *U. unicinctus* exhibits strong sulfide tolerance by enhancing the sugar metabolism pathway and increasing its energy production to compensate for the lack of aerobic respiration capacity caused by repetitive sulfide stress.

## 5. Conclusion

*U. unicinctus* enhances plasticity by triggering a sulfide stress memory mechanism to enable acclimation. In the present study, we describe the phenomenon of sulfide stress memory in organisms by establishing a sulfide stress memory system in *U. unicinctus*. Further, we obtained TMGs involved in stress memory by analyzing transcriptomic data, classified them according to their expression patterns, and functionally analyzed them. Ultimately, we obtained the regulatory network of key pathways and genes involved in sulfide stress memory of *U. unicinctus*. In response to repeated sulfide stress, *U. unicinctus* enhances its cellular protection by enhanced responses of TMGs in pathways such as sulfide metabolism, antioxidant enzyme systems, energy metabolism, and maintenance of protein balance. Additionally, the modified response of TMGs involved in DNA repair may result in decreased sensitivity to repeated sulfide stress. This study provides new insights into the mechanisms of adaptation to sulfide stress in organisms. Although we successfully established the sulfide stress memory system of *U. unicinctus*, other concentrations that can induce sulfide stress memory and the required recovery time need to be further tested. Furthermore, the duration of sulfide stress memory and whether it is heritable in *U. unicinctus* are currently unknown. The molecular mechanisms regulating sulfide stress memory and the functions of important TMGs also require further exploration.

## Credit authorship contribution statement

**Wenqing Zhang:** Writing – original draft, Visualization, Methodology, Investigation. **Danwen Liu:** Visualization, Methodology, Investigation, Data curation. **Heran Yang:** Methodology, Data curation. **Tianya Yang:** Methodology, Data curation. **Zhifeng Zhang:** Supervision, Conceptualization. **Yubin Ma:** Writing – review & editing, Supervision, Funding acquisition, Conceptualization.

## Data availability statement

The transcriptome data presented in this study are deposited in the NCBI repository with the accession number PRJNA752504 and PRJNA1112745.

## Acknowledgment

This work was supported by the PhD Scientific Research and Innovation Foundation of Sanya Yazhou Bay Science and Technology City (HSPHDSRF-2022-02-010), Shandong Province Science Outstanding Youth Fund (ZR2020YQ20), China Postdoctoral Science Foundation (2020M680095), the Fundamental Research Funds for the Central Universities and Qingdao Postdoctoral Application Research Project.

## Supporting Information

**Figure A.1.** Schematic diagram of sulfide treatment and sampling for *U. unicinctus*.

**Figure A.2.** Pearson correlation analysis and PCA analysis of sequencing samples. Pearson correlation analysis of sequencing samples. (b) PCA analysis of sequencing samples.

**Figure A.3.** Transcription data and qRT-PCR analysis of TMGs in sulfide metabolism.

(a) Transcription data and qRT-PCR analysis of *sqr*. (b) Transcription data and qRT-PCR analysis of *aox*. (c) Transcription data and qRT-PCR analysis of *sod*. (d) Transcription data and qRT-PCR analysis of *cat*. Different letters indicate significant differences between the data (*P* <0.05).

**Figure A.4.** Heat map of expression levels of key enzyme genes (*sqr*, *tst*, and *pdo*) in sulfide metabolism pathway under repeated sulfide stress.

**Table A.1.** qRT-PCR primer list.

**Table A.2.** RNA sequencing data and corresponding quality control.

**Table A.3.** List of reads and reference genome comparison.

**Table A.4.** Expression patterns of TMGs at various points in time and typology.

**Table A.5.** Top10 hub genes in module 2.

**Table A.6.** Top10 hub genes in module 12.

**Table A.7.** Top10 hub genes in module 8.

