## Supplementary figures and images for "Transcriptional memories mediate the plasticity of sulfide stress responses to enable acclimation in *Urechis unicinctus*"

### Figure A1

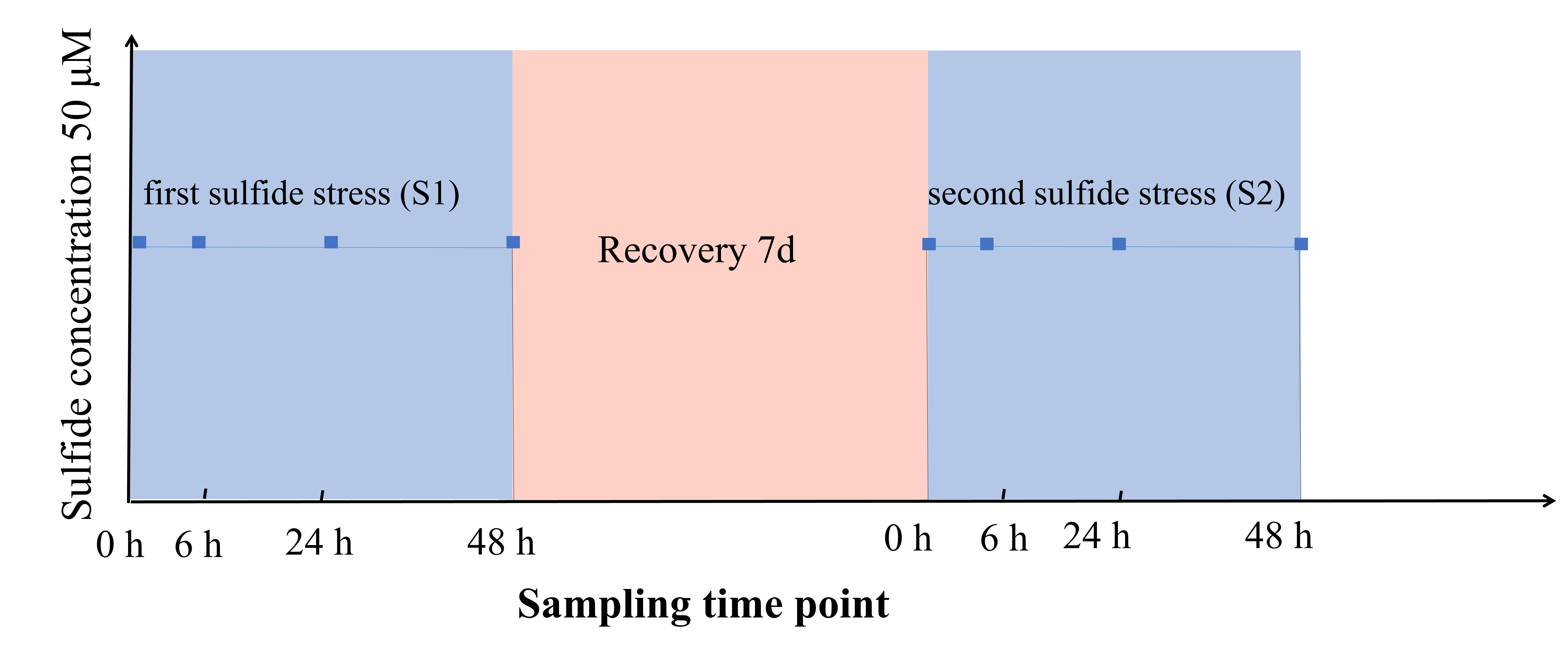

### Figure A2

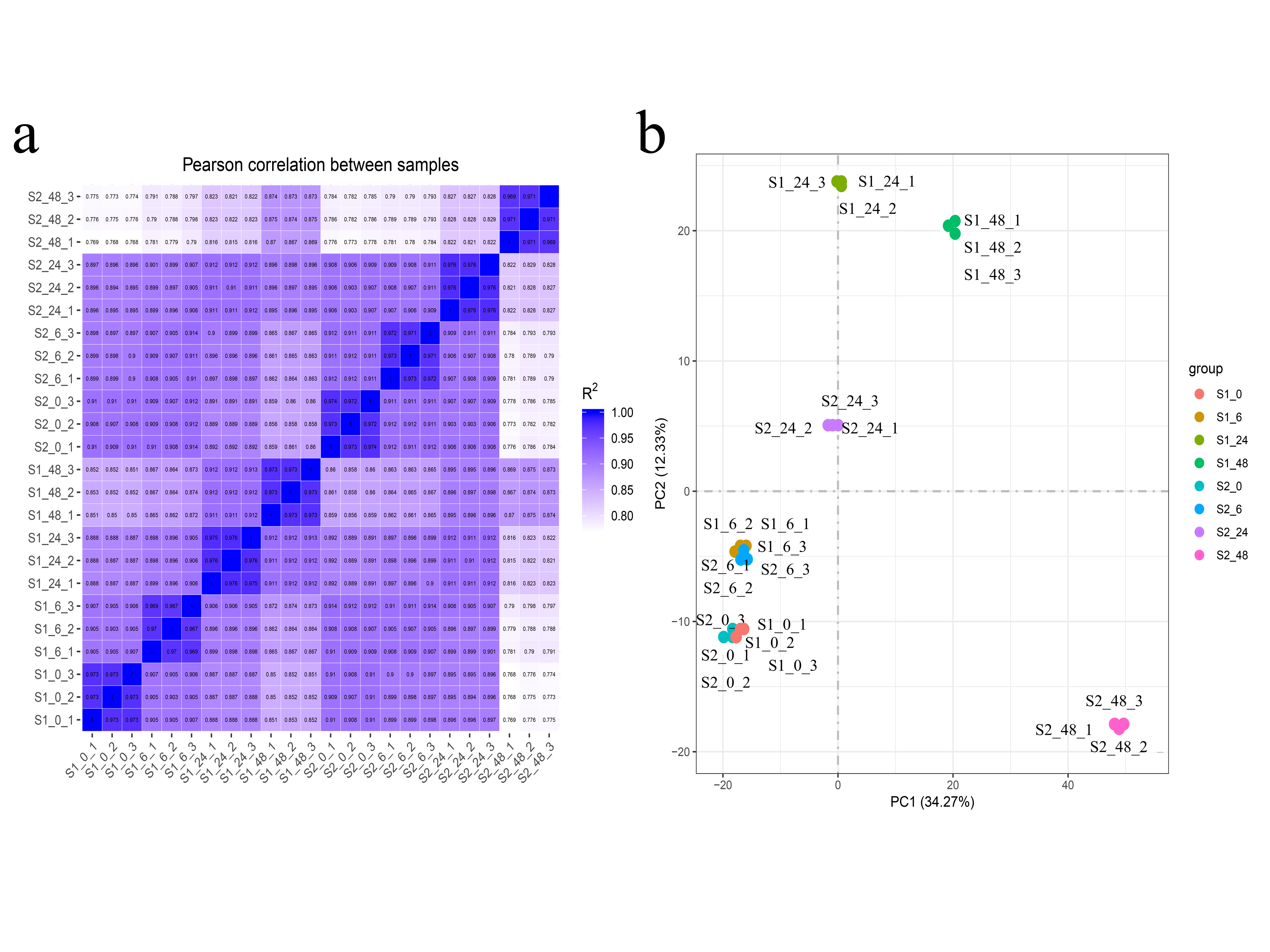

### Figure A3

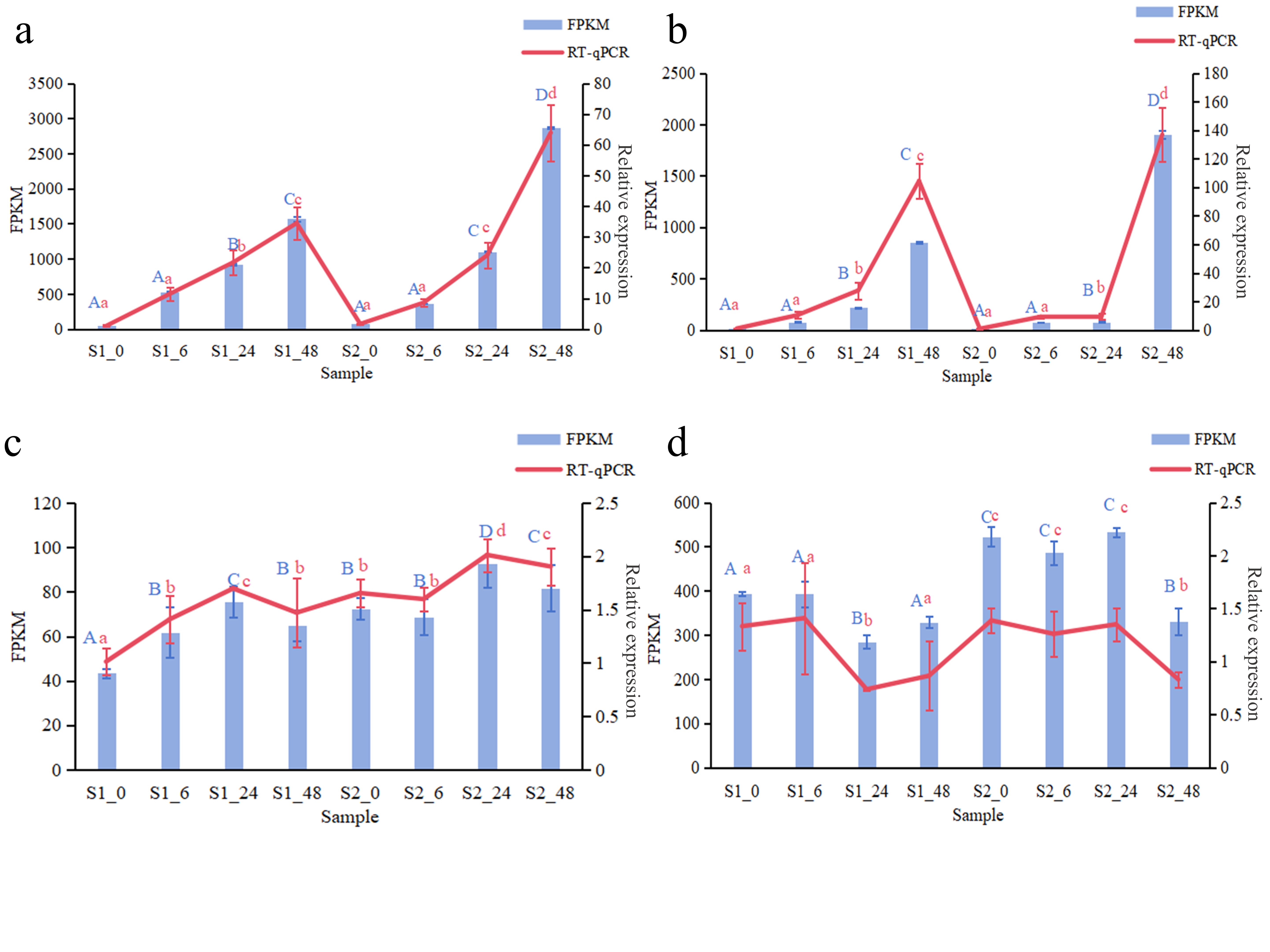

### Figure A4

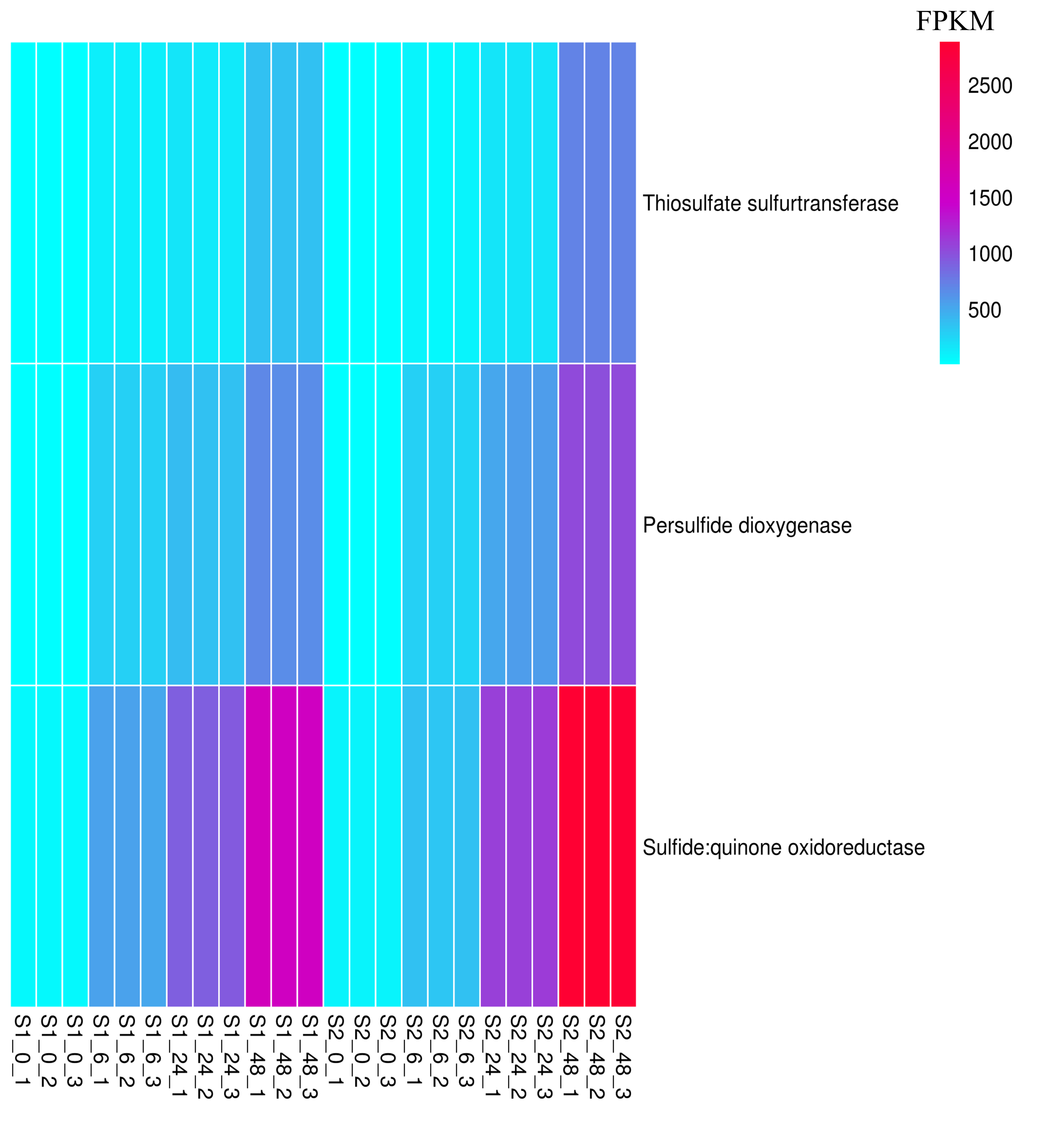
